# Nutrient Concentration Declines but Nutrient Yield Increases Under Elevated CO_2_ Concentration in Soybean: Disentangling the relative effects of Dilution, Transpiration, and Uptake Activity

**DOI:** 10.64898/2026.08.28.747630

**Authors:** Terence Seldon Kwafo, Kylie Yerkes, Justin M. McGrath

## Abstract

Rising atmospheric CO_2_ concentrations consistently reduce mineral concentrations in C_3_ crops, yet the mechanisms driving this decline – dilution, reduced transpiration-driven mass flow, and altered root nutrient acquisition capacity – have rarely been tested simultaneously under field conditions. Here, we grew two soybean (Glycine max Merr.) varieties with contrasting yield responses to elevated [CO_2_] (Loda and HS93-4118) at the Soybean Free Air CO_2_ Enrichment (SoyFACE) facility and quantified the independent contributions of dilution, transpiration, and root uptake to observed changes in tissue mineral concentration and nutrient yield. Elevated [CO_2_] increased grain yield by 21% and 6% in Loda and HS, respectively, and reduced transpiration by 13-14% across both varieties. A decomposition analysis revealed that while dilution and reduced transpiration each exerted negative effects on tissue nutrient concentrations, with their combined effect representing a 31- 40% potential reduction in whole-plant concentration, active root uptake responses were positive for nearly all nutrients, largely offsetting these losses and resulting in observed concentration declines substantially smaller than either mechanism alone would predict. Consequently, whole-plant nutrient yield increased under elevated CO_2_ for all elements measured, and grain nutrient yield increased for most. The relationship between seasonal transpiration and nutrient yield was steeper under elevated CO_2_ for most macronutrients, indicating that plants acquired more nutrients per unit water transpired, not less, under elevated CO_2_. These results demonstrate that root uptake is the dominant compensatory response to elevated [CO_2_] driven dilution and reduced mass flow in soybean, that nutrient yield is maintained or increased for most elements under elevated [CO_2_], and that Fe represents a physiologically and nutritionally significant vulnerability in future CO_2_ environments.

## Introduction

Atmospheric CO_2_ concentrations ([CO_2_]) (elevated [CO_2_]) influence crop nutrient composition, with potential implications for both crop yield and quality, with deficiencies in any essential element significantly reducing growth and crop yields (Marschner, 1995). Changes in nutrient concentrations in major food crops may also affect nutritional quality of harvested products, consequently reducing human dietary mineral intake (Loladze, 2014; Myers *et al*., 2014a). Mineral nutrients are essential for plant growth and human nutrition. Despite this dependence, mineral malnutrition remains a major health challenge: approximately two-thirds of the world’s population is deficient in one or more essential dietary nutrients (Bird *et al*., 2017). Iron (Fe) and zinc (Zn) deficiencies are widespread public health concerns, linked to impaired immunity, inadequate physical development, and increased susceptibility to infection (WHO, Micronutrient deficiencies, 2020).

Elevated [CO_2_] affects plant growth and development by increasing photosynthetic rate but also by reducing stomatal conductance (Woodward, 2002; Ainsworth and Long, 2005; Leakey *et al*., 2009). Enhanced photosynthesis can increase biomass production, grain yield, and water-use efficiency in C_3_ grasses and legumes (Ainsworth *et al*., 2002, 2008; Leakey *et al*., 2012). However, despite improved plant growth under elevated [CO_2_], grain quality is often reduced in many crops (Loladze, 2014; Myers *et al*., 2014a). Free-air CO_2_ enrichment (FACE) studies have shown that plants grown under elevated [CO_2_] often have lower nutrient concentrations than plants grown under ambient conditions (Loladze, 2014; Taub et al., 2008). In rice, wheat, and soybean, elevated [CO₂] has been associated with concentration reductions of 2- 20% for nutrients including nitrogen (N), potassium (K), calcium (Ca), magnesium (Mg), Fe, Zn, manganese (Mn), and sulphur (S) (Manderscheid *et al*., 1995; Högy and Fangmeier, 2008; Fernando *et al*., 2014; Myers *et al*., 2014b; Guo *et al*., 2015).

Although many studies show that elevated [CO_2_] changes crop quality, the mechanisms underlying declines in plant mineral concentrations remain unresolved. Three hypotheses are commonly mentioned, though the strength of evidence supporting each varies considerably. The first hypothesis is nutrient dilution, whereby increased photosynthetic carbon assimilation raises non-structural carbohydrate levels and reduces mineral concentrations if nutrient uptake does not increase proportionally (Poorter *et al*., 1997). Under this hypothesis, lower nutrient concentrations may not necessarily indicate reduced mineral uptake, only a change in the ratio between carbon gain and mineral uptake.

A second hypothesis proposes that reduced stomatal conductance under elevated [CO_2_] lowers transpiration, thereby decreasing mass-flow driven nutrient transport to roots. Because this process depends on nutrient solubility in soil water, it affects highly soluble nutrients more than relatively insoluble nutrients. For example, a nutrient such as Fe, which is relatively insoluble, would be transported at a lower concentration in the soil solution than a highly soluble nutrient such as Ca. Therefore, unlike the dilution hypothesis, the transpiration hypothesis predicts that nutrient responses to elevated [CO_2_] should differ according to nutrient solubility. Several studies have failed to support the transpiration hypothesis (Muenscher, 1922; Wong, 1979; Tanner and Beevers, 1990; Conroy, 1992; Rudmann *et al*., 2001). In contrast, McGrath and Lobell (2013) leveraged the expected differential responses of highly soluble versus less soluble nutrients and reported support for the hypothesis, an approach not used in earlier studies. However, this meta-analysis paired CO_2_-induced nutrient concentration changes across studies with literature-derived estimates of soil nutrient mobility rather than with direct within-experiment measurements, an approach that may introduce artifacts since nutrient solubility varies with soil type. Transpiration rates themselves were not measured directly, and most data came from chamber or greenhouse studies in which differences in canopy coupling to the atmosphere would affect transpiration rates.

A third hypothesis is that elevated [CO_2_] may alter root-system nutrient-acquisition capacity. Nutrient accumulation depends not only on the movement of nutrients through the soil solution, but also on the capacity of roots to absorb, transport, and allocate those nutrients to tissues. Elevated [CO_2_] can stimulate belowground biomass allocation and fine root proliferation, potentially increasing the surface area available for nutrient acquisition (Pritchard and Rogers, 2000; Taub and Wang, 2008*b*; Tausz-Posch *et al*., 2014). However, increased root biomass does not necessarily translate into proportionally greater nutrient uptake. Some evidence also suggests that root functional capacity for nutrient uptake may be suppressed under elevated [CO_2_] with no increase in root biomass (BassiriRad *et al*., 2001; Kaur *et al*., 2025). Changes in root architecture, transporter activity, or root metabolic activity under elevated [CO_2_] can decouple morphological expansion from functional nutrient absorption capacity (Taub and Wang, 2008; Beidler *et al*., 2015; Jauregui *et al*., 2016). Additionally, soil nutrient availability is determined by soil chemical properties, physical structure, and plant nutrient acquisition strategies, which complicates efforts to disentangle the relative contributions of dilution, transpiration-driven mass flow, and altered uptake capacity to nutrient declines under elevated [CO_2_].

Because nutrient concentrations often decline in plants grown under elevated [CO_2_], it is commonly assumed that the net effect of rising [CO_2_] on human nutrition will be detrimental. However, this assumption does not account for the yield stimulation frequently observed under elevated [CO_2_], which could potentially offset reduced nutrient concentrations by increasing the total amount of nutrients produced per unit land area (Farhoomand and Peterson, 1968; Digrado *et al*., 2024). Whether the net effect is beneficial or harmful depends on the relative magnitudes of nutrient concentration declines and yield increases, as well as broader effects on prices, purchasing choices, and dietary composition. Although this study cannot address all those downstream factors, it can directly test whether nutrient yield, defined as the mass of nutrient produced per unit ground area, is altered by growth under elevated [CO_2_]. If nutrient yield decreases, then reduced nutrient concentrations would likely translate into lower nutritional output, as is often assumed. However, if nutrient yield is maintained or increased, then assessing whether growth in elevated [CO_2_] necessarily worsens nutritional outcomes becomes less straightforward.

Disentangling the relative contributions of these hypotheses requires experiments that directly measure transpiration, tissue nutrient concentrations, nutrient accumulation, biomass production, and nutrient yield under realistic field conditions. Previous studies have shown that soybean varieties (cultivars) differ considerably in their yield responses to elevated [CO_2_] (Bishop *et al*., 2015), providing an opportunity to test whether nutrient responses vary with growth stimulation. Therefore, this study used a FACE facility to evaluate two soybean varieties, Loda and HS93-4118, with contrasting yield responses to elevated [CO_2_]. Because Loda has been shown to exhibit a greater yield increase than HS93-4118 under elevated [CO_2_] (Sanz-Sáez *et al*., 2017), these varieties were used to examine the relative effects of the proposed mechanisms. The objective was to determine the degree to which reductions in soybean nutrient concentration under elevated [CO_2_] are explained by each of dilution, reduced transpiration-driven mass flow, altered nutrient acquisition capacity. We tested two hypotheses: **(1)** the relative contribution of dilution, reduced transpiration, and altered nutrient acquisition capacity differs among nutrients and soybean varieties under elevated [CO_2_]; and **(2)** growth under elevated [CO_2_] reduces nutrient yield. Together, these analyses evaluate whether elevated [CO_2_] reduces soybean nutritional quality through changes in concentration alone, or whether changes in biomass production and nutrient yield modify the net nutritional consequences of rising atmospheric [CO_2_].

## Materials and Methods

### Study site and experimental setup

Field studies of soybean plants in ambient and elevated [CO_2_] using a FACE system were conducted during the 2022-2024 growing seasons. To elevate atmospheric CO_2_ concentrations, plants were grown in plots inside a 20 m diameter octagonal ring of pipes to fumigate the plot with CO_2_ (**Fig. 1**), hereafter called rings. The plants inside each ring were compared with an adjacent plot of plants grown outside the ring as a control group. To control for environmental variation, the experiment was repeated across several pairs of rings and adjacent control plants in each year in a randomized complete-block design with each pair as a block. There were four ring pairs in 2022 and 2023, but only three in 2024. Two soybean varieties, Loda (PI 614088) and HS93-4118 (PI 614155), were planted in each ring with a plot of four 2 m long rows with a row spacing of 0.76 m (30 in). Loda was planted all three years, while HS93-4118 was only planted in 2023 and 2024. The [CO_2_] treatment for ambient (control) plots was 415 ppm, and elevated fumigated with CO_2_ to reach a target of 600 ppm. Experimental details of CO_2_ fumigation are described by Aspray *et al*. (2023). The elevated CO_2_ rings were fumigated during the day throughout the growing season from emergence to maturity (Miglietta *et al*., 2001).

**Fig. 1.**
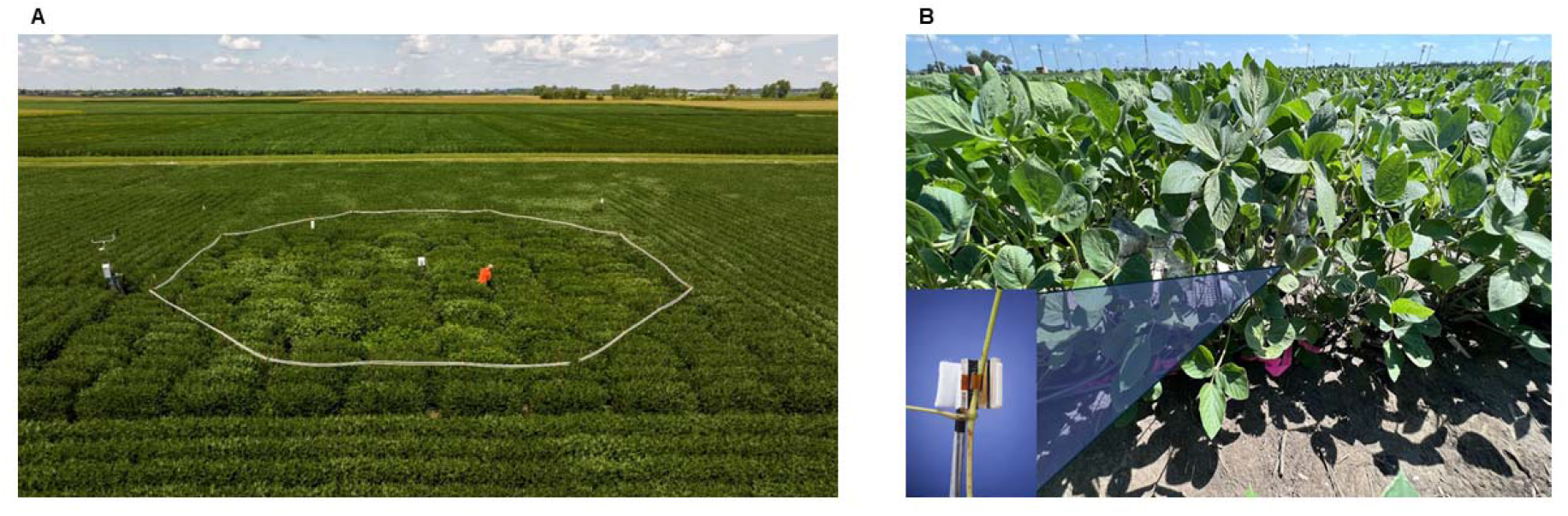
Experimental plots at SoyFACE field site. **(A)** Aerial picture of one of the experimental plots, where plants were exposed to ambient or elevated [CO_2_]. Elevated [CO_2_] ring pictured has section of white pipes where CO_2_ is released on the upwind sides of the plots. **(B)** Soybean plants with sap flow gages installed.

The study was conducted as at the Soybean Free Air Concentration Enrichment (SoyFACE) facility in Champaign, IL, USA (40°02′ N, 88°14′ W, 228 m above sea level; Rogers *et al*., 2004). SoyFACE is a 32-ha farmland with soybean (*Glycine max* [L.] Merr.) and maize (*Zea mays*) planted in rotation and managed according to common practices (Bishop *et al*., 2015). The farm is a rainfed site with tile drainage with Drummer silty clay loam and Flanagan silt loam soil types. Weather information was obtained from the weather station located at the SoyFACE facility (**Fig. 2**).

**Fig. 2.**
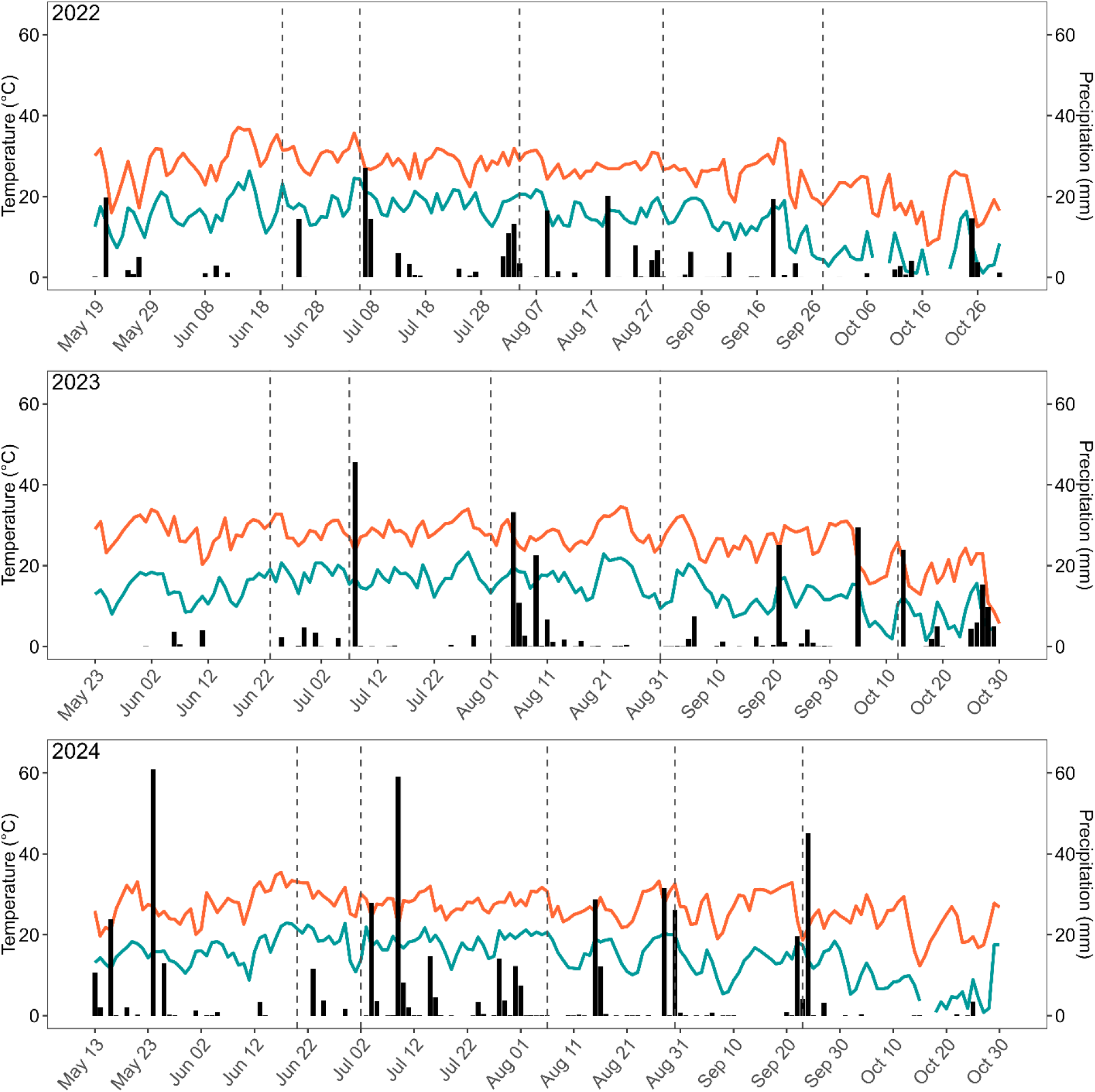
Daily weather conditions during the growing season from planting date to harvest in 2022 - 2024. Precipitation is represented as black vertical bars. Maximum and minimum temperatures are indicated by the solid orange and blue lines, respectively. Data were obtained from the SoyFACE weather station. Dash vertical lines denote sampling dates.

### Gas exchange

Leaf-level gas exchange measurements were taken after canopy closure at R5 on two consecutive days to parameterize the Ball et al. (1987) gs model. Two randomly selected mature leaves from each plot were cut from the canopy before dawn. Petioles were recut underwater and then kept in low-light conditions before taking measurements (Ainsworth et al., 2006). Measurements were performed on a mature central leaflet using a gas exchange system with a 6 cm^2^ leaf chamber (LI-6800; Li-Cor, Lincoln, NE, USA). Conditions within the instrument were as follows: ambient [CO_2_] = 415 μmol mol^−1^, elevated [CO_2_] = 600 μmol mol^−1^, flow = 500 μmol m^-2^ s^-1^, fan speed = 1,000 RPM, temperature (T) = 27 °C, relative humidity (RH) = 70%. Photosynthetic photon flux density (PPFD) incident on the leaf was varied stepwise (2000, 1200, 800, 600, 400, 300, 200 μmol m^−2^ s^−1^). Data were recorded for each stepwise change in PPFD when A and gs reached steady state, which required 10-30 minutes per point. This methodology was modified from Leakey et al. (2006). The stomatal conductance (gs), intercept (g0), and slope (m) of the Ball et al. (1987) model were estimated using the calculate_ball_berry_index function from the PhotoGEA R package, which fits light response curves (Lochocki et al., 2025).

### Plant tissue collection, soil sampling and nutrient analysis

To measure aboveground biomass yield and nutrient yield throughout the season, a 2-meter section of a row was randomly selected in each plot and harvested at five growing stages: vegetative stage 3 (V3), beginning bloom (R1), beginning seed (R5), full seed (R6), and full maturity (R8) (Fehr and Caviness, 1977) (**Table S1**). For grain yield, a 2 m row was harvested from each plot at full maturity. Within the area sampled 3-6 plants were partitioned into leaves, stems, roots, and pods to estimate tissue fractions and for nutrient analysis. Samples were dried for two weeks at 60 °C in a drying oven before weighing.

The concentration of total macro (N, P, K, S, Ca, and Mg) and micro (Fe, Mn, Zn, Cu, B, Al, and Na) nutrients in tissue samples were analysed by inductively coupled plasma atomic emission spectrometry (ICP-AES) at a service laboratory (Waypoint Analytical, Champaign, IL). Soil core samples 0.02 m in diameter and 0.25 m deep were collected from the plots. Two subsamples were pooled, one from the center and one from the side near the perimeter, from each plot.

Nutrient yield was calculated as nutrient concentrations multiplied by biomass yield expressed per unit ground area (Equation 2.1).

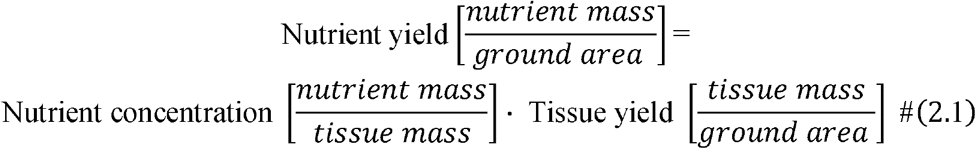

### Sap flow and transpiration estimates

Transpiration was measured on four whole soybean plants per variety in each ring using a heat balance method (Sakuratani, 1981) with sap flow sensors attached to the lower third of the stem so that water flux through the plant would be measured (SGA5-WS, Dynamax Inc., Houston, TX, USA). The sensors were installed around the stems between two nodes when the stem diameter reached between 5 - 7 mm (**Fig. 1B**). To minimize stem temperature variation, which would interfere with measurements, the sensors were covered with insulation foam. A second aluminium bubble foil layer was secured over the foam to shield from radiant loading, prevent spikes in sensor reading, and reduce condensation. Installation was performed according to the instrument manual (Dynagage Sap Flow Sensor - Dynamax Inc., Houston, TX, USA). Each week plants and sensors were checked for damage. The sap flow measurements were recorded using a data logger (CR1000X, Campbell Scientific Inc., Logan, UT, USA) with a proprietary program provided by the manufacturer (Dynamax Inc., Houston, TX, USA).

Sap flow rates (g h^-1^) were recorded at 15-minute intervals for ∼40 days. The means of those values over the growing season were compared between treatments to determine effect of elevated [CO_2_] on transpiration. Sap flow rates were summed to daily totals (g day^-1^) for each calendar day, then the daily rate per plant, *F* (g day^-1^ plant^-1^), was converted to transpiration per plot, *T* (mm day^-1^ m^-2^), using the planting density, *ρ_d_* = 39.3701 plants m^-2^, the density of water, *ρw* = 1,000 g L^−1^ and Equation (2.2).

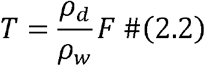

Daily transpiration rates *T* were then averaged over the season for each year (August 7 - September 10 in 2022; July 25 - September 10 in 2023 and 2024) to obtain a seasonal daily mean transpiration (mm day^-1^).

### Nutrient uptake mechanism in response to Elevated [CO_2_]

We analyzed the relative impacts of dilution, transpiration, and increased root mass on the observed change in tissue nutrient concentration under elevated CO_2_ as follows.

Nutrient yield is not impacted by dilution; only concentration changes due to the increase in carbohydrate mass. If dilution were the only effect – that is, if absolute nutrient yield remained unchanged between treatments while biomass increased by a factor of (1 + r) – then the expected change in concentration under elevated CO_2_ would be

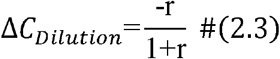

where *r* is the relative increase in biomass yield (B) under elevated CO_2_ 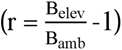. The dilution term is strictly negative whenever CO_2_ stimulates biomass accumulation, reflecting the expectation that nutrient concentrations decline proportionally as fixed nutrient pools are distributed across a larger biomass (**Equation S1**).

To calculate the expected transpiration effect, we calculated the hypothetical nutrient yield (*Y_hyp_*) under elevated [CO_2_], assuming that nutrient delivery remained determined solely by the ambient treatment relationship between transpiration and nutrient yield. For each treatment, the yield *Y_i_* of nutrient *i* (g m^-2^) was first regressed against cumulative transpiration *T* (mm) for each nutrient and tissue type using ordinary least squares:

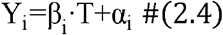

where β_i_ is the slope, and α_i_ is the y-intercept. Regressions were fitted separately for ambient and elevated CO_2_ treatments. Both varieties, Loda and HS, had similar linear relationships within each CO_2_ treatment with no evidence of variety-specific deviation. Y_hyp_ was then computed as the predicted nutrient yield at the observed elevated [CO_2_] transpiration rate using the soil water nutrient concentration and the reduction in transpiration between treatments:

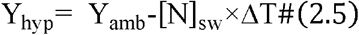

where [N]_SW_ is the nutrient concentration in soil water (**Table S3B**), and ΔT is the difference in transpiration between treatments (ΔT = T_amb_-T_elev_).

The difference between Y_hyp_ and the ambient treatment’s predicted yield at ambient transpiration isolates the contribution of reduced transpirational mass flow to changes in nutrient supply. Independent of any change in root uptake capacity, this term estimates the effect of reduced transpiration:

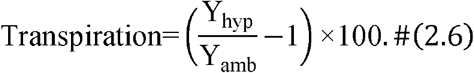

The combined effect is the non-additive interaction between dilution and the transpiration-driven reduction in nutrient supply, that is, the degree to which simultaneous increases in biomass and reductions in transpiration jointly change tissue concentration:

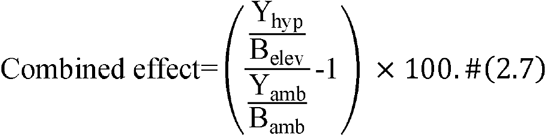

The difference between the observed elevated CO_2_ yield (Y_elev_) and Y_hyp_ captures the residual attributable to another response, uptake activity.

Hence, the uptake activity effect represents the residual change in nutrient concentration that could be attributed to active physiological adjustments in nutrient acquisition, including changes in root architecture, transporter expression or activity not accounted for by the dilution and transpiration driven mass-flow mechanisms above:

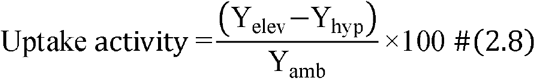

where a positive uptake activity term indicates that plants acquired more nutrients relative to ambient.

### Statistical analysis

Statistical analysis was performed on plot means. When subsamples were collected, the mean of subsamples within a plot was used. For all analyses, there were four replicates (n=4) for 2022 and 2023 and three replicates (n=3) for 2024. Because results from field experiments are often highly variable, a false positive rate of 10 % was chosen (α = 0.1). A mixed model analysis of variance (ANOVA) type III split-plot design as carried out using the lmerTest R package to obtain the p-values (Kuznetsova *et al*., 2017). The general model structure:

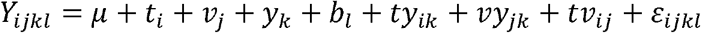

where *Y_ijkl_* is the response variable; μ is the overall mean; (*t_i_*) and (*v_j_*) are fixed effects; (*y_k_*), *block* (*b_l_*), (*ty_ik_*), and (*vy_jk_*) random effects; (*tv_ij_*) is the fixed interaction term; and *ε_ijkl_* is the residual error. Interaction terms in the model that were not significant were removed.

## Results

### Meteorological and climatic conditions for duration of experiment

Growing season cumulative precipitation amounts for the three consecutive years were 235 mm, 229 mm, and 475 mm (**Fig. 2; Table S2**). The years 2022 and 2023 were the driest, while 2024 was the wettest. The average maximum air temperature was consistent across all years at 27 ℃. Mean maximum VPD was 1.4 kPa in each of 2022 and 2023, higher than the observed 1.17 kPa in 2024 (**Fig. S1; Table S1**).

### Grain yield increased under elevated [CO_2_]

Grain yield response to elevated [CO_2_] differed significantly between varieties and across the three growing seasons (**Fig. 3**). Averaged across years, Loda and HS yields increased by 21% and 6% under elevated [CO_2_], respectively (*p* < 0.005). Loda responded four times more strongly than HS, despite HS maintaining greater total above-ground biomass throughout the experiment (**Fig. S2**), suggesting that the two varieties differed in their capacity to partition elevated [CO_2_]-driven carbon gain into grain production. Harvest index was not significantly affected by elevated [CO_2_] in either variety (T: *p* < 0.898; **Fig. S2**). Loda had a significantly greater harvest index than HS (V: *p* < 0.074; **Fig. S2**).

**Fig. 3.**
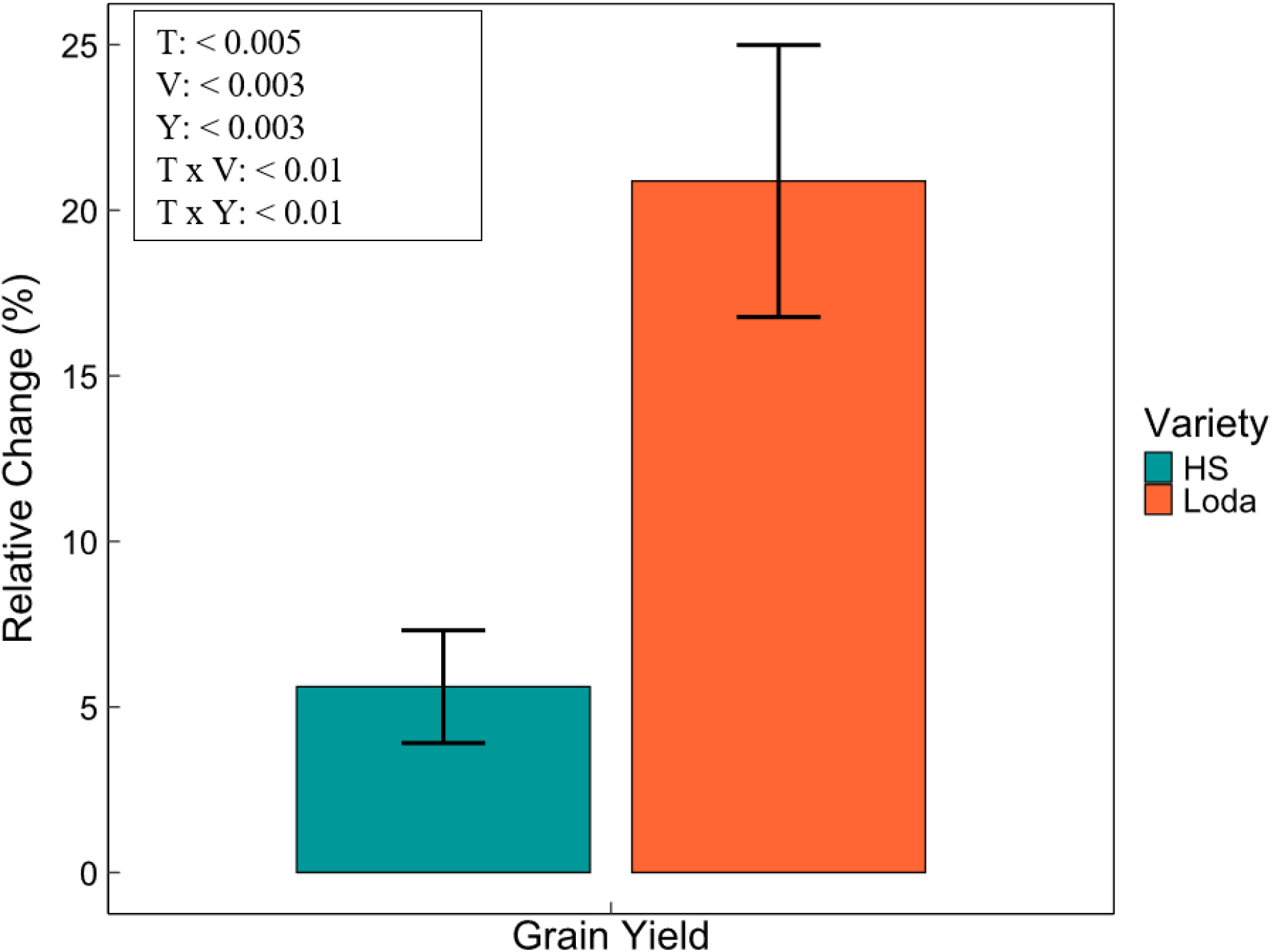
**Change (%) in yield in response to elevated CO_2_ relative to ambient levels**. HS93-4118 (HS) and Loda are represented in blue and orange, respectively. Bars are relative change in the mean yield between the ambient and the elevated CO_2_ treatments: (E-A)/A, where A and E are the mean yield of the ambient and the elevated CO_2_ treatments respectively. Error bars represent 95% CI. The impacts of various factors on yield were analyzed using a mixed linear model: Yield ∼ Treatment (T) + Variety (V) + Year (Y). A Type III ANOVA was then performed to assess statistical significance, with *p*-values for the final model given in the inset.

### Elevated [CO_2_] reduces seasonal transpiration

Daily transpiration averaged 6.9 mm day^-1^ under ambient conditions and 5.9 mm day^-1^ under elevated [CO_2_] across both varieties and all years (**Fig. 4A**). Average daily transpiration was significantly different across years and between CO_2_ treatments, while varieties were not significantly different (treatment: *p* = 0.004; year: *p* < 0.001; variety: *p* = 0.71). Transpiration rate varied among years of the experiment, with 2024 being the highest and 2022 the lowest water use for both varieties. Across all three years and both varieties, growth under elevated [CO_2_] consistently resulted in lower seasonal cumulative transpiration compared to ambient [CO_2_] (**Table 1**). Elevated [CO_2_] reduced seasonal transpiration in both cultivars, with a three-year mean reduction of 13.9% for Loda and a two-year mean reduction of 12.8% for HS (**Table 1**). The lower stomatal conductance in plants grown under elevated [CO_2_] supported the transpiration responses to elevated [CO_2_]. Across years and varieties, elevated [CO_2_] reduced stomatal conductance relative to ambient conditions, with mean values declining by 29% for HS and 14% for Loda (T: p < 0.06; V: p < 0.048; Y: p < 0.004; **Fig. 4B**).

**Fig. 4.**
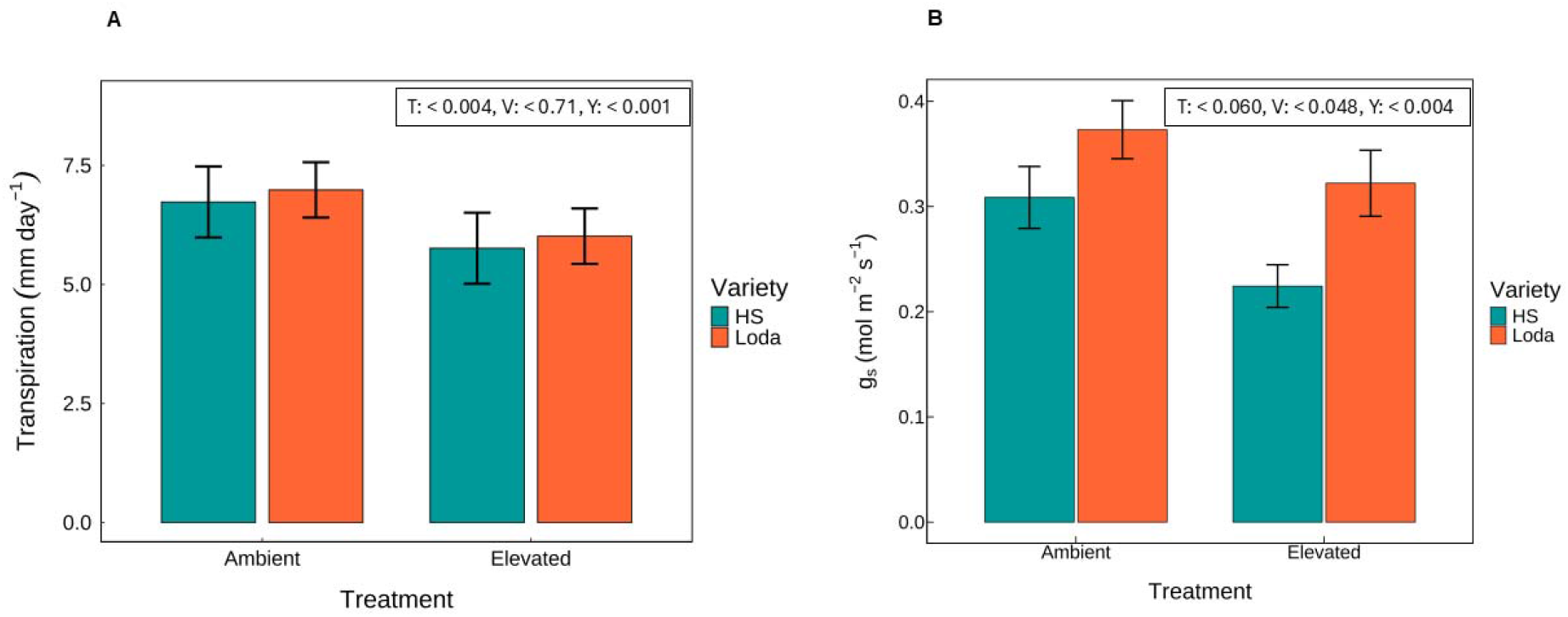
(A) Average daily transpiration rate (mm day^-1^). (B) Mean stomata conductance (g_s_) (mol m^−2^ s^−1^). HS93-4118 (HS) and Loda are represented in blue and orange, respectively. Error bars represent 95% CI. The data were analyzed using a mixed linear model: Transpiration rate/Stomatal conductance ∼ Treatment (T) + Variety (V) + Year (Y). A Type III ANOVA was then performed to assess statistical significance, with *p*-values for the final model given in the inset.

**Table 1.** Cumulative transpiration (mm season^-1^) from 2022 to 2024.

| Year | Variety | Ambient | Elevated | % Change |
| --- | --- | --- | --- | --- |
| 2022 | Loda<br>HS | 440.3 | 358.6 | -18.5 |
| 2023 | Loda<br>HS | 622.8<br>601.6 | 541.2<br>520.0 | -13.1<br>-13.6 |
| 2024 | Loda<br>HS | 697.1<br>675.9 | 615.5<br>594.2 | -11.7<br>-12.1 |
| **Mean** | **Loda**<br>**HS** | **586.7**<br>**638.7** | **505.1**<br>**557.1** | **-13.9**<br>**-12.8** |

Values shown are three or four replicate blocks, depending on the year. Measurements (mm season<sup>-1</sup>) are from the sensor installation date until senescence.

### Elevated [CO_2_] reduces nutrient concentration but increases nutrient yield

#### Whole plant nutrient concentration

Elevated [CO_2_] broadly reduced whole-plant (aboveground tissues; sum of leaves, stems, and pods) and grain nutrient concentrations relative to ambient levels at seed filling (R5) and full maturity (R8) for both varieties (**Fig. 5**). Nutrient concentrations differed by variety, treatment and years (**Fig. 5A**; **Table 2**). For whole-plant measurements, elevated [CO_2_] decreased N concentration (*p* = 0.002), with a significant treatment-by-variety interaction (*p* = 0.004; *p* < 0.1 in all cases; **Table 2B**). Whole-plant K, Ca, and Mn concentrations also tended to decline under elevated [CO_2_] (K, *p* = 0.039; Ca, *p* = 0.080; Mn, *p* = 0.062), whereas P and Mg were unaffected by CO_2_ but varied between varieties (P, *p* = 0.077; Mg, *p* = 0.097; **Table 2B**), consistent with the low soil water P (**Table S3B**). HS had a greater decline for most nutrients except for K and Cu. Fe concentration had the largest mean decline of 19%. Zn concentration also decreased under elevated [CO_2_], although inconsistently between varieties.

**Fig. 5.**
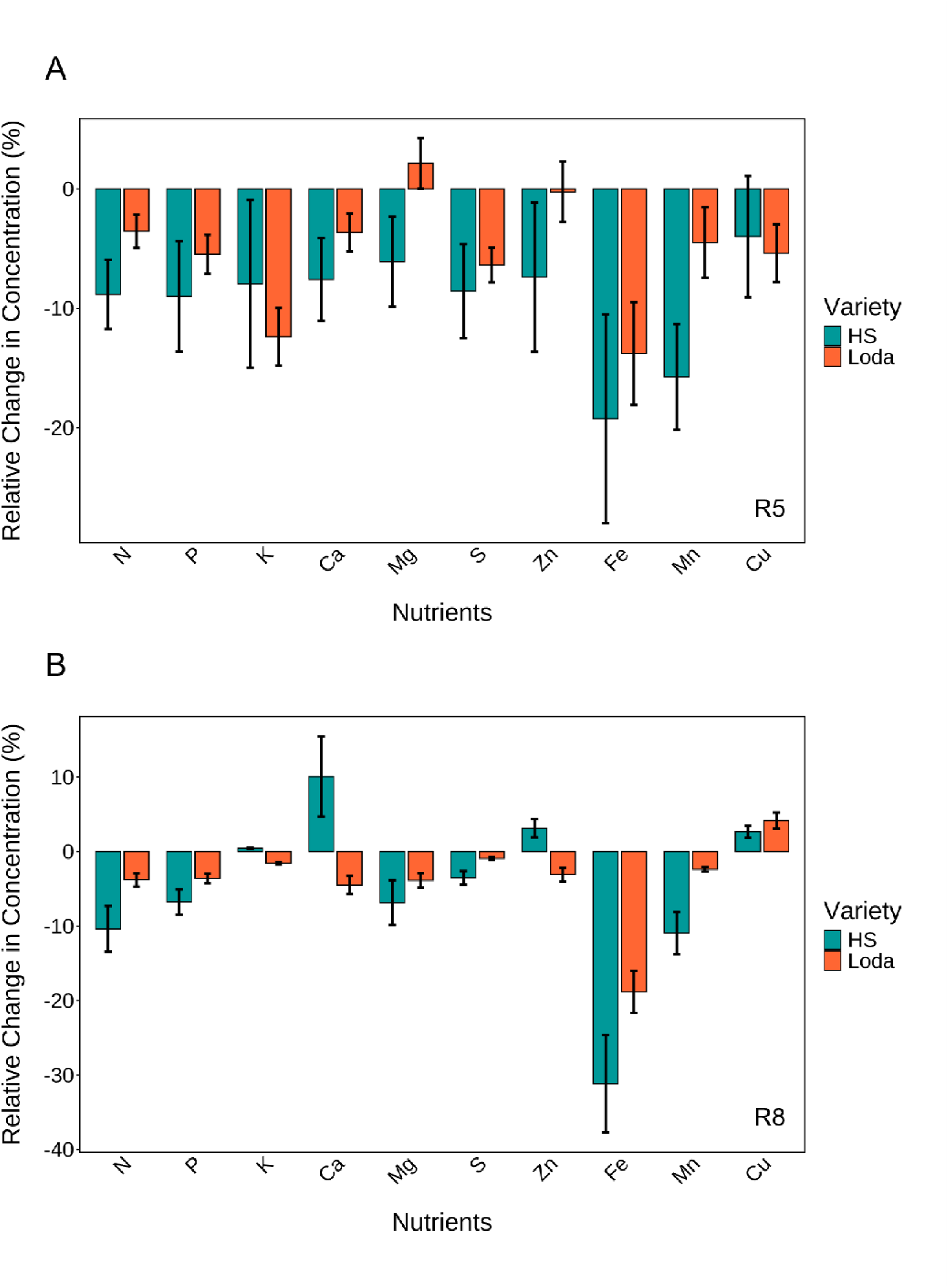
**Nutrient concentration change (%) in response to elevated CO_2_ relative to ambient levels**. **(A)** Whole plant (aboveground) nutrient concentration at development stage R5**. (B)** Grain nutrient concentration at development stage R8. HS93-4118 (HS) and Loda are represented in blue and orange, respectively. Error bars represent 95% CI. The relative change in the mean concentration between the ambient and the elevated CO_2_ treatments: (E-A)/A, where A and E are the mean yield of an at the ambient and the elevated CO_2_ treatments respectively. The impact of various factors on nutrient concentration ([N]) was analyzed using a mixed linear model: [N] ∼ Treatment (T) + Variety (V) + Year (Y). A Type III ANOVA was then performed to assess statistical significance with *p-values* for the final model.

**Table 2.** ANOVA of nutrient concentration in soybean grain at R8 and whole plant (aboveground) at R5.

| **(A) Grain** |  |  |  |  |  |  |
| --- | --- | --- | --- | --- | --- | --- |
| **Nutrient** | **CO<sub>2</sub> Treatment (T)** | **Variety (V)** | **Year (Y)** | **T x V** | **T x Y** | **V x Y** |
| **N** | 0.194 | 0.644 | **0.036** | 0.632 | 0.416 | 0.189 |
| **P** | 0.458 | 0.139 | **0.055** | 0.690 | 0.510 | 0.570 |
| **K** | 0.614 | 0.547 | 0.117 | 0.778 | 0.381 | 0.540 |
| **Ca** | 0.741 | 0.150 | **0.079** | **0.400** | 0.756 | 0.337 |
| **Mg** | 0.549 | 0.337 | **0.035** | **0.473** | 0.957 | 0.528 |
| **S** | 0.488 | 0.954 | **0.053** | **0.617** | 0.571 | 0.272 |
| **Zn** | 0.923 | 0.933 | **0.058** | **0.806** | 0.626 | 0.355 |
| **Fe** | **0.057** | 0.519 | 0.198 | 0.391 | 0.391 | 0.579 |
| **Mn** | 0.270 | 0.307 | **0.082** | 0.386 | 0.725 | 0.228 |
| **Cu** | **0.036** | **0.050** | **0.003** | 0.115 | **0.039** | **0.012** |
| **(B) Whole plant** |  |  |  |  |  |  |
| **Nutrient** | **CO<sub>2</sub> Treatment (T)** | **Variety (V)** | **Year (Y)** | **T x V** | **T x Y** | **V x Y** |
| **N** | **0.002** | **0.003** | **0.006** | **0.004** | **0.004** | **0.007** |
| **P** | 0.126 | **0.077** | 0.546 | 0.504 | 0.255 | 0.303 |
| **K** | **0.039** | **0.083** | 0.144 | 0.163 | 0.239 | 0.800 |
| **Ca** | **0.080** | 0.112 | **0.087** | 0.216 | 0.216 | **0.093** |
| **Mg** | 0.323 | **0.097** | 0.111 | 0.112 | 0.796 | 0.332 |
| **S** | **<0.001** | **<0.001** | **<0.001** | **<0.001** | **<0.001** | **<0.001** |
| **Zn** | 0.351 | 0.117 | **0.045** | 0.320 | 0.359 | 0.679 |
| **Fe** | 0.325 | 0.191 | 0.233 | 0.647 | 0.533 | 0.147 |
| **Mn** | **0.062** | **0.077** | **0.022** | **0.092** | 0.155 | 0.266 |
| **Cu** | 0.110 | 0.251 | 0.044 | 0.540 | 0.411 | 0.258 |

*p*-values are presented for the main effects of treatment (T), variety (V), and year (Y), as well as for all two- and three-way interactions affecting seed elemental concentrations. Non-significant two- or three-way interactions were excluded, and the model was refitted. Bold *p*-values indicate significant differences at alpha = 0.1

#### Grain nutrient concentration

At R8, grain nutrient concentrations were generally lower in elevated [CO_2_], though not consistently significantly different (**Fig. 5B**; **Table 2A**). N, P, K, Mg and S grain concentration declines not significant. Grain Fe and Cu declined significantly (Fe, *p* = 0.057; Cu, *p* = 0.036), with a 30% reduction for Fe. Zn showed a marginal decline of 5% only for Loda. The pronounced grain Fe concentration decline coincided with consistently low soil water Fe availability across all years (**Table S3B**), likely impacted by the soil pH range of 6.1–6.8 (**Table S3A**), which may have reduced Fe^3+^ solubility and constrained Fe loading into the grain.

#### Nutrient yield responses to elevated [CO_2_]

Aboveground tissue nutrient yields increased in all cases in elevated [CO_2_] despite the clear reductions in concentrations (**Fig. 6A**; **Table 3B**). HS generally had a larger increase in nutrient yield for all nutrients except Mn. In contrast, several grain nutrient yields increased for most nutrients, except Fe, which decreased (**Fig. 6B**; **Table 3A**).

**Fig. 6.**
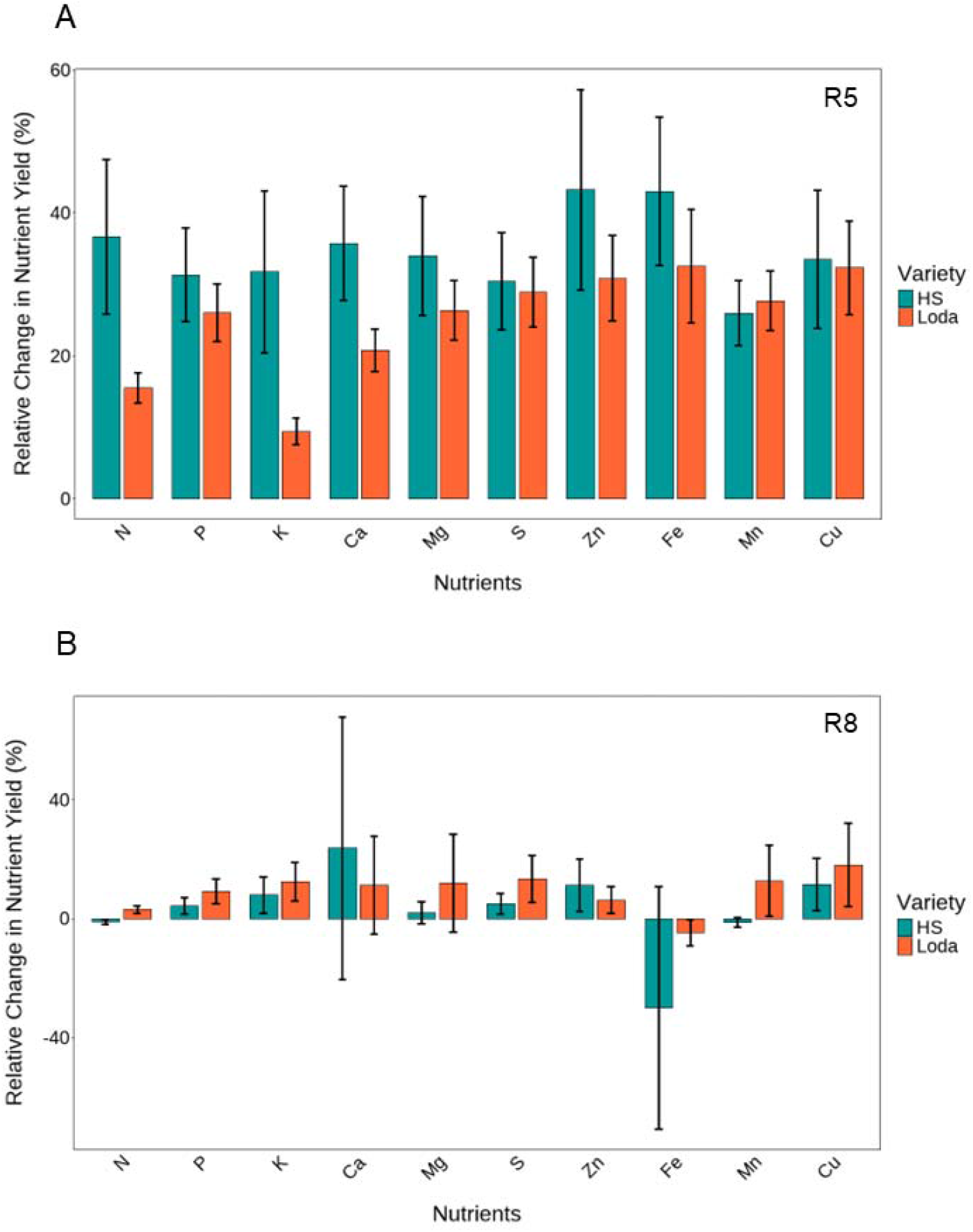
Nutrient yield change (%) in response to elevated CO_2_ relative to ambient levels. **(A)** Whole plant (aboveground) nutrient yield at development stage R5**. (B)** Grain nutrient yield at development stage R8. HS93-4118 (HS) and Loda are represented in blue and orange, respectively. Error bars represent 95% CI. The relative change in the mean concentration between the ambient and the elevated CO_2_ treatments: (E-A)/A, where A and E are the mean yield of an at the ambient and the elevated CO_2_ treatments respectively. The impact of various factors on nutrient yield was analyzed using a mixed linear model: Nutrient yield ∼ Treatment (T) + Variety (V) + Year (Y), along with their interactions. A Type III ANOVA was then performed to assess statistical significance with *p-values* for the final model.

**Table 3.** ANOVA of nutrient yield in soybean grain at R8 and whole plant (aboveground) at R5.

| **(A) Grain** |  |  |  |  |  |  |
| --- | --- | --- | --- | --- | --- | --- |
| **Nutrient** | **CO<sub>2</sub> Treatment (T)** | **Variety (V)** | **Year (Y)** | **T x V** | **T x Y** | **V x Y** |
| **N** | 0.850 | 0.463 | 0.289 | 0.807 | 0.919 | 0.444 |
| **P** | 0.435 | 0.207 | 0.463 | 0.703 | 0.650 | 0.824 |
| **K** | 0.184 | 0.289 | 0.140 | 0.529 | 0.871 | 0.706 |
| **Ca** | 0.131 | 0.115 | **0.033** | 0.300 | 0.211 | 0.250 |
| **Mg** | **0.029** | 0.602 | **0.006** | 0.263 | **0.052** | 0.978 |
| **S** | 0.170 | 0.184 | 0.219 | 0.329 | 0.732 | 0.215 |
| **Zn** | 0.477 | 0.537 | 0.200 | 0.910 | 0.858 | 0.497 |
| **Fe** | 0.291 | 0.997 | 0.147 | 0.375 | 0.622 | 0.948 |
| **Mn** | **0.002** | **0.068** | **0.000** | **0.003** | **0.005** | **0.003** |
| **Cu** | **0.004** | **0.005** | **0.004** | **0.015** | **0.018** | **0.005** |
| **(B) Whole plant** |  |  |  |  |  |  |
| **Nutrient** | **CO<sub>2</sub> Treatment (T)** | **Variety (V)** | **Year (Y)** | **T x V** | **T x Y** | **V x Y** |
| **N** | 0.178 | 0.629 | 0.155 | 0.482 | 0.997 | 0.257 |
| **P** | 0.270 | 0.272 | 0.153 | 0.733 | 0.781 | 0.245 |
| **K** | 0.411 | 0.628 | 0.195 | 0.412 | 0.891 | 0.292 |
| **Ca** | 0.233 | 0.758 | 0.373 | 0.676 | 0.896 | 0.289 |
| **Mg** | 0.178 | 0.300 | 0.320 | 0.793 | 0.859 | 0.497 |
| **S** | 0.194 | 0.473 | 0.152 | 0.458 | 0.859 | 0.265 |
| **Zn** | 0.214 | 0.343 | 0.297 | 0.509 | 0.642 | 0.326 |
| **Fe** | 0.740 | 0.340 | 0.218 | 0.632 | 0.751 | 0.456 |
| **Mn** | 0.164 | 0.602 | **0.053** | 0.535 | 0.460 | 0.137 |
| **Cu** | 0.162 | 0.596 | 0.202 | 0.372 | 0.810 | 0.398 |

*p*-values are presented for the main effects of treatment (T), variety (V), and year (Y), as well as for all two- and three-way interactions affecting seed elemental concentrations. Non-significant two- or three-way interactions were excluded, and the model was refitted. Bold *p*-values indicate significant differences at alpha = 0.1

#### Interaction between nutrient yield and transpiration

Plants grown in elevated [CO_2_] accumulated more nutrients per volume of water transpired (**Fig. 7)**. Nutrient-yield–transpiration relationship slopes were steeper under elevated [CO_2_] for the majority of nutrients. For nitrogen, the slope increased by 17% under elevated [CO_2_] (0.1068 g m^-2^ mm^-1^). Slopes also increased for P (0.0084 g m^-2^ mm^-1^), K (0.036 g m^-2^ mm^-1^), Ca (0.052 g m^-2^ mm^-1^), and Mg (0.022 g m^-2^ mm^-1^). The steeper slopes for Ca and Mg are consistent with their relatively high concentrations in soil water (**Table S3B**). Slopes for Mn and Zn were unchanged between treatments, while Fe was the only nutrient for which the slope was slightly lower under elevated [CO_2_]. This trend was similar for grains at final harvest at R8 (**Fig. S3**). On average, nutrient yield per the amount of water transpired was 16% and 29% higher (N, P, K, Ca, Mg) under elevated [CO_2_], for whole-plant and grain, respectively. Both varieties, HS and Loda, had consistent nutrient uptake with no evident genotypic differences in the transpiration–nutrient relationship (**Fig. 7**).

**Fig. 7.**
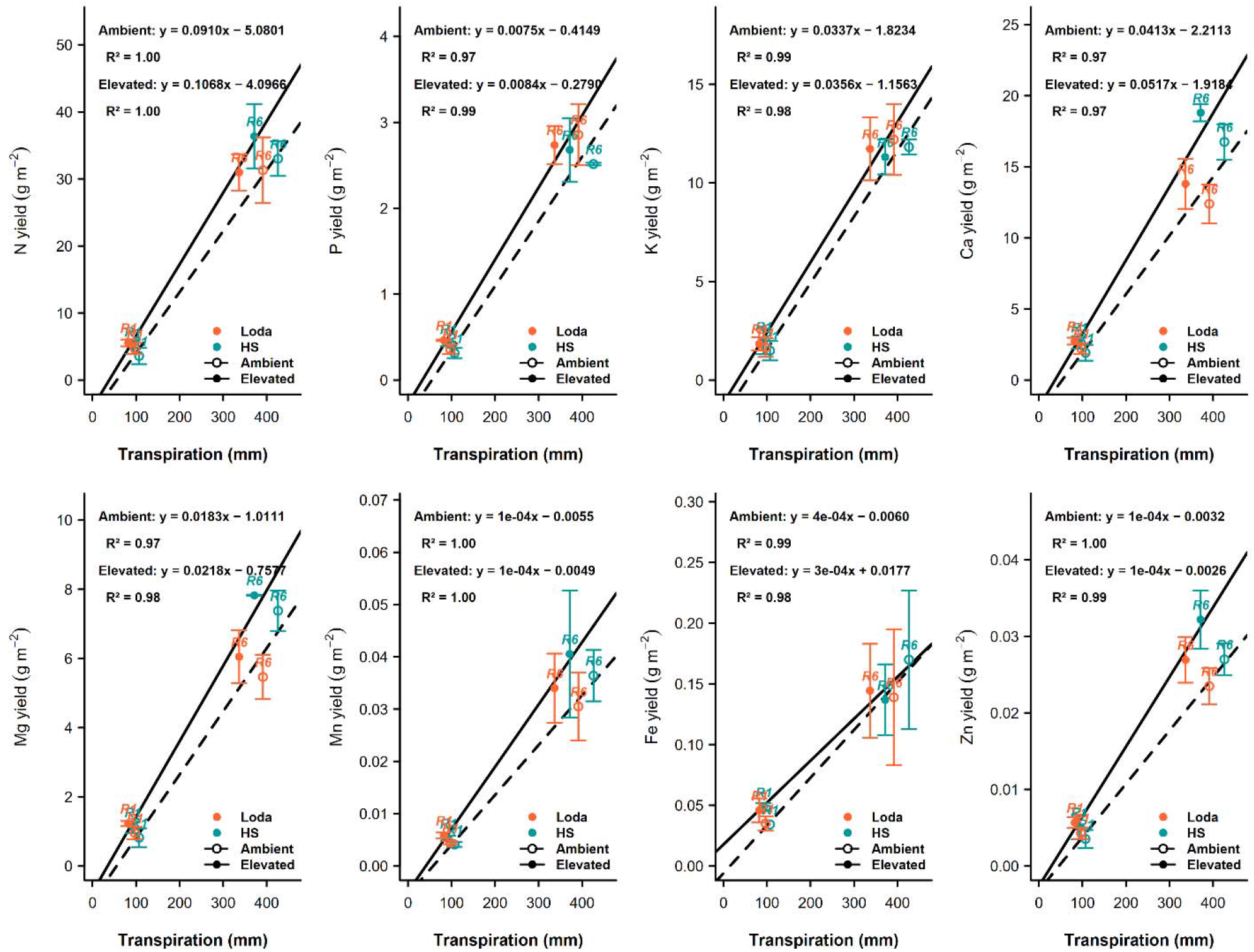
Effects of transpiration on nutrient uptake in response to elevated CO_2_. Whole plant (aboveground) nutrient response at development stage R5. HS93-4118 (HS) and Loda are represented in blue and orange, respectively. Ambient CO_2_ (dashed lines and open circles) and Elevated CO_2_ (bold lines and filled circles). Error bars represent 95% CI.

#### Relative effects of mechanism contribution to nutrient concentration decline in responses to elevated [CO_2_]

The observed concentration changes represent the measured nutrient responses to elevated [CO_2_], whereas the dilution and transpiration effects represent the expected contributions to those responses. Uptake activity was inferred as the difference between the observed and expected responses; positive values indicate increased acquisition or allocation that offset these negative effects.

### Dilution effect

In whole-plant aboveground tissue, elevated [CO_2_] generally reduced the observed nutrient concentrations of both varieties. The measured declines were greater in HS than in Loda for most nutrients, except K and Cu. This pattern corresponded with the stronger whole-plant biomass response of HS and, consequently, its larger dilution effect. Pure dilution was estimated to reduce nutrient concentrations by 31.0% in HS (r = 0.45), compared with 20.0% in Loda (r = 0.25) (**Fig. 2.8A–B**). Dilution was also a larger effect than reduced transpiration for every nutrient in both varieties. These results suggest that biomass dilution was a major contributor to the stronger whole-plant nutrient declines in HS. However, the observed declines in whole-plant concentration were substantially smaller than those expected from the combined effects of dilution and transpiration. Despite dilution effects of −31.0% in HS and −20.0% in Loda, most observed declines remained between 0% and −20%. The relationship between dilution and the observed response differed in grain because the varieties showed contrasting grain-yield responses. Loda had the greater grain-yield response to elevated [CO_2_], and consequently the larger expected grain dilution effect, at −16.7% (r = 0.20), compared with only −5.7% in HS (r = 0.06) (**Fig. 2.8C–D**). The small dilution effect in HS was consistent with its smaller grain-yield response. In HS grain, reduced transpiration had a larger expected effect than dilution for most nutrients. In contrast, dilution was larger than the transpiration effect for most nutrients in Loda grain, except Ca, Mg, and Zn. These differences indicate that the relative importance of dilution depends on the grain-yield response of each variety. The stronger grain-yield stimulation in Loda increased the contribution of dilution.

### Transpiration effect

The expected transpiration effect varied by nutrient availability and solubility in soil water. Ca and Mg showed the largest transpiration effects in grain for HS and Loda (**Fig. 2.8C–D; Table 3**), consistent with relatively high soil water Ca and Mg concentrations (**Table 3B**). Nevertheless, the observed grain concentration changes were much smaller in most cases than these expected declines. For P, K, Mn, Fe, and generally Zn, the transpiration effect was smaller due to their lower soil water concentrations. Grain Fe showed a pronounced observed decline in uptake activity, particularly in HS. Unlike the other nutrients, Fe acquisition or allocation to the grain did not increase sufficiently to offset dilution and reduced transpiration.

Overall, the observed concentrations changes were only partially explained by dilution and transpiration. Dilution was most influential where elevated [CO_2_] produced the strongest biomass response in whole-plant HS and grain Loda. Reduced transpiration contributed to a lesser and more selective degree, limited to nutrients present at relatively high concentrations in soil water. For most nutrients, however, the observed declines were smaller than the expected concentration decline from these two mechanisms alone indicating that increased root uptake activity provided substantial compensation.

## Discussion

This study aimed to parse the relative contributions of dilution, reduced transpiration-driven mass flow, and altered root uptake or acquisition capacity to decline in mineral concentrations observed in soybean grown under elevated [CO_2_]. The decomposition analysis of these hypotheses revealed that while both dilution and reduced transpiration exert negative effects on tissue nutrient concentrations, increased uptake activity largely offset their combined impact for most nutrients, resulting in observed concentration declines that were substantially smaller than dilution and transpiration mechanisms would predict (**Fig. 8**). Moreover, despite consistent declines in concentration, whole-plant and grain nutrient yield increased under elevated [CO_2_] across nearly all elements (**Fig. 6**), complicating the widely held assumption that rising atmospheric CO_2_ will necessarily reduce the nutritional output of soybean crops.

**Fig. 8.**
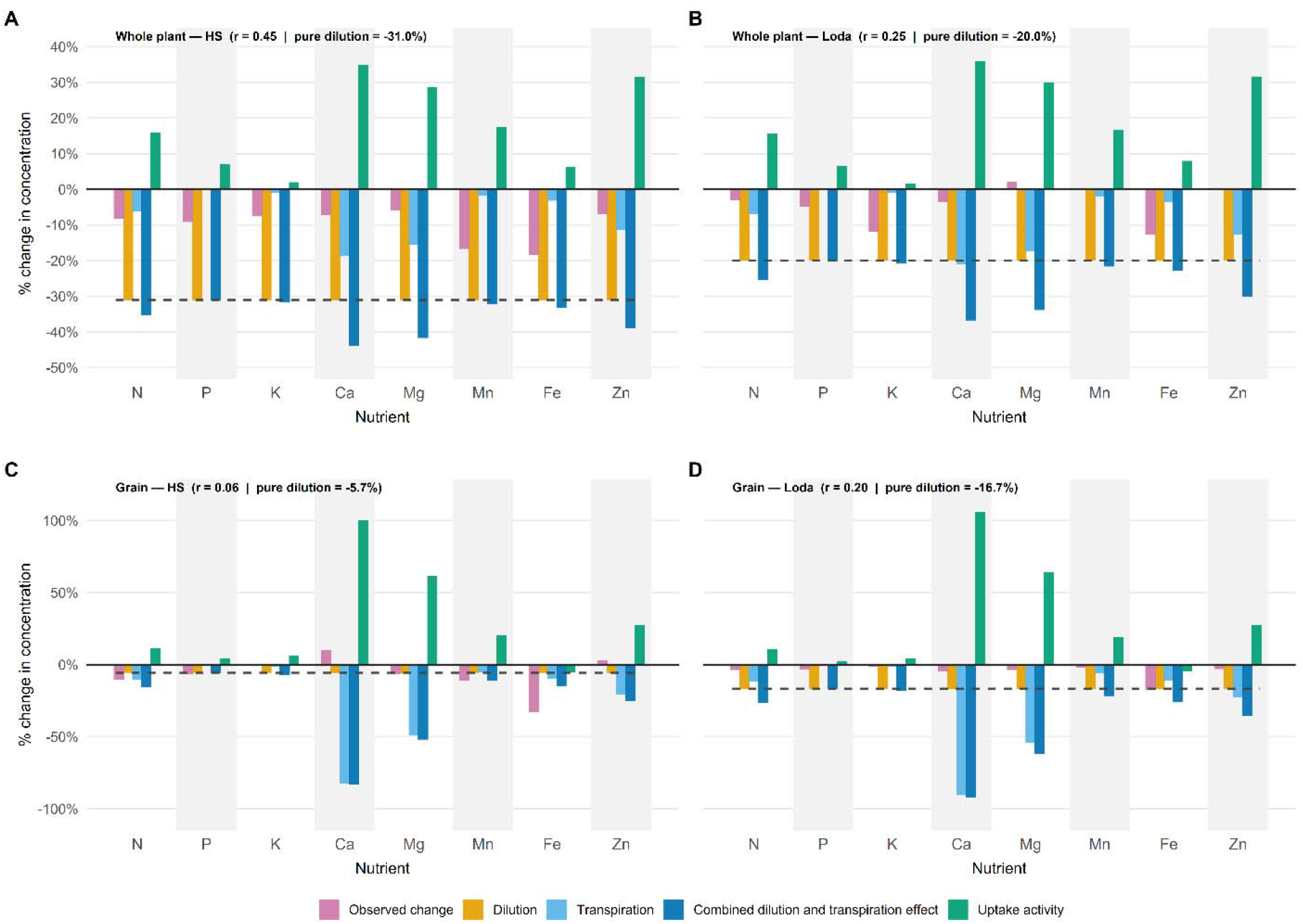
Relative effects of nutrient decline mechanisms. (A-B) Effects on HS and Loda, whole plant (aboveground) at seed fill (R5). (C - D) Effects on HS and Loda, grain at final harvest (R8). r is the relative increase in biomass yield under elevated CO_2_. Pink for observed concentrations; Light blue for transpiration; Dark blue for combined transpiration and dilution effect; Orange bar for dilution; and Green for positive and negative root uptake, respectively.

### Dilution occurs but does not act alone

The expected dilution effect is uniform across nutrients within a given tissue and variety, since it depends solely on the ratio of carbon gain to mineral accumulation (**Fig. 8**), meaning that all non-photosynthetically assimilated elements are affected equally. The predicted magnitude scaled directly with biomass stimulation under elevated [CO_2_], and differed between tissue types and varieties, reflecting the well-established principle that elevated [CO_2_] does not stimulate all organs equally (Poorter *et al*., 1997; Ainsworth and Long, 2005). In aboveground tissue, where biomass stimulation was greater for HS but lower for Loda, the predicted dilution effect exceeded that in grain (**Fig. 3; Fig. S2**), consistent with the minimal harvest index response of HS compared to Loda in response to elevated [CO_2_]. However, observed concentration changes were consistently larger than expected from dilution alone for most nutrients in aboveground tissue for both varieties. In some cases, for instance in HS grain, the observed change in nutrient concentration was more negative than the dilution effect (**Fig. 2.8 C**), indicating that dilution alone cannot explain the observed nutrient decline; additional mechanisms contributed to concentration declines beyond what dilution imposed. These patterns confirm that while dilution consistently reduces tissue nutrient concentration under elevated [CO_2_], the magnitude of the observed concentration change ultimately depends on whether active root uptake compensates for that loss, a finding with important implications for modeling future crop nutritional quality.

### Minimal effects due to transpiration are offset or mitigated by other physiological mechanisms

Transpiration and stomatal conductance were significantly reduced under elevated [CO_2_] across both varieties and all three growing seasons (**Fig. 4**), as observed previously (Bernacchi *et al*., 2006, 2007; Myers *et al*., 2014a; Kaur *et al*., 2025), and this reduction was sustained across dry and wet years (**Fig. 4**; **Table 1, S1**). The potential magnitude of the transpiration effect on nutrient concentration scales directly proportional to soil water nutrient concentration (**Table S3**), and thus differed substantially between nutrients and tissue types. The hypothetical transpiration effect on grain Ca and Mg concentrations, which had the highest soil water concentrations, substantially exceeded the dilution effect in both varieties (**Fig. 8C & 8D**). For N, P, K, Mn, Fe, and Zn, where soil water concentrations were considerably lower, the transpiration effect was smaller than dilution (**Fig. 8**; **Table 3**). However, observed concentration changes did not follow that pattern, indicating that active root uptake responses for these nutrients were strongly positive across both varieties, nearly entirely offsetting the losses.

The relationships between transpiration and whole-plant or grain nutrient yield were also steeper under elevated than ambient [CO_2_] for most macronutrients, including N, P, K, Ca, and Mg (**Fig. 7; Fig. S3**). Plants therefore accumulated more nutrient per unit of water transpired under elevated [CO_2_], consistent with Houshmandfar *et al*. (2018). This finding does not support reduced transpiration as a major limitation to total nutrient acquisition. Instead, it indicates that nutrient uptake, transport, or allocation increased sufficiently to compensate for lower water flux. The similar responses of Loda and HS, despite differences in biomass and grain-yield stimulation, suggest that this compensation was not strongly variety specific. Root responses provide a plausible mechanism for this compensation. Elevated [CO_2_] frequently increases root length, diameter, branching, rooting depth, and the number of root tips, although responses vary among species and may be restricted by nutrient limitation (Rascher *et al*., 2010; Cohen *et al*., 2018, 2019; Uddin *et al*., 2018). Both HS and Loda had larger root yield under elevated [CO_2_] (**Fig. S2A**). A larger or deeper root system could increase the volume of soil explored and maintain nutrient acquisition despite lower mass-flow delivery. However, greater root growth under elevated [CO_2_] can also be accompanied by a 20–40% reduction in root xylem area (Cohen *et al*., 2018, 2019). Thus, increased soil exploration and reduced hydraulic transport may occur simultaneously: greater absorptive surface could support nutrient uptake, while reduced xylem capacity reinforces lower transpiration and limits passive nutrient transport to the shoot.

Soybean nutrient demand also remains high during reproductive development, where N, P, Ca, Mg, S, Zn, Mn, B, and Cu acquisition is distributed between vegetative growth and seed filling, rather than being completed early in development (Bender *et al*., 2015). <u>Gaspar *et al*., (2018)</u> similarly reported that 48–73% of secondary and micronutrient uptake occurred at flowering and seed filling. These patterns suggest that the positive uptake-activity (**Fig. 8**) may reflect continued root acquisition during reproductive growth, when elevated [CO_2_]-stimulated biomass and grain development increased nutrient demand.

### Altered uptake capacity and transporter activity may compensate for most nutrients

Nutrient accumulation depends not only on transport through the soil solution, but also on root uptake, radial transport, xylem loading, remobilization, and allocation to developing tissues. For most nutrients, the declines predicted from dilution and reduced transpiration were substantially greater than the observed changes in concentration (**Fig. 8**). Thus, these two mechanisms alone could not explain the measured response. Within the decomposition framework, the remaining difference is attributed to altered uptake activity, indicating that nutrient acquisition or allocation increased under elevated [CO_2_].

Most nutrients showed evidence of this physiological compensation. In most cases, there was positive uptake activity for N, P, K, Mn, and Zn for whole-plant and grain, with whole-plant having a greater increase (**Fig. 8**). Compensation was especially strong for Ca and Mg. Although reduced transpiration predicted larger expected declines in their grain concentrations, the observed responses were small. These patterns suggest that increased root acquisition, transport, or allocation largely maintained nutrient supply despite lower transpirational mass flow. Greater root surface area, continued uptake during seed filling, and increased transporter activity could all contribute to this response. It is also consistent with the induction of high-affinity membrane transporters when nutrient demand increases (Marschner, 2008).

Fe concentration declined strongly, particularly in grain, and Fe was the only nutrient for which grain nutrient yield decreased under elevated [CO_2_] (**Fig. 5**; **Fig. 6**). For Fe, dilution and reduced transpiration accounted for most of the observed decline, leaving little or no additional contribution attributed to altered uptake activity (**Fig. 7**; **Fig. 8**). This result is particularly important because soybean grain [Fe] depends strongly on continued uptake from the soil. Gaspar et al. (2018) found that Fe accumulated in soybean seed after R5 came from direct soil uptake rather than remobilization from vegetative tissues. Fe uptake also remained relatively high through maturity. Consequently, limitations to Fe solubilization, root uptake, radial transport, xylem loading, or movement into developing grain during seed filling would directly restrict grain Fe accumulation. This Fe-specific response is physiologically plausible because Fe availability is strongly constrained by soil chemistry (Lindsay and Schwab, 1982; Marschner, 1995; Saleem *et al*., 2023). Although Fe is abundant in many soils, especially at the SoyFACE field site (**Table S3A**), it is often present in oxidized, poorly soluble forms that are not readily available for plant uptake.

As a dicot, soybean uses Strategy I Fe acquisition, a sequential process of rhizosphere acidification by plasma membrane H^+^-ATPases (AHA2), ferric reductase activity (FRO2) to reduce Fe^3+^ to the soluble Fe^2+^ form, and subsequent transport through transporters such as IRT1 (Marschner, 1995; Molnár *et al*., 2023). IRT1 is known to mediate the uptake of multiple divalent metal cations including Fe, Zn, Mn, and Cu (Vert *et al*., 2002; Marschner, 2008; Connorton *et al*., 2017). The broadly positive active uptake response observed for Zn, Mn, and Cu in this study may therefore partly reflect coordinated upregulation of IRT1 activity driven by increased nutrient demand under elevated [CO_2_] stimulated growth. Where Zn, Mn and Cu act as structural components of proteins, enzyme activators, and key regulators of photosynthesis and stress responses. However, the limited supply of soluble Fe may have prevented a comparable compensatory response for Fe. Evidence from rice also indicates that elevated [CO_2_] can reduce the expression of OsZIP3 and OsZIP5, which participate in the transport of Fe, Zn, Mn, and Cu (Ainsworth *et al*., 2025). Although these responses have not been established in soybean, they demonstrate that elevated [CO_2_] may affect grain micronutrient concentration through transporter regulation in addition to dilution and transpiration. The Strategy I pathway is also critically sensitive to soil pH: FRO2-mediated reduction activity declines sharply above pH 6.5–7.0, and at the near-neutral to slightly alkaline pH characteristic of the SoyFACE site soils, Fe^3+^ solubility is inherently limited (**Table S3A**)(Lindsay and Schwab, 1982; Marschner, 1995). Under these conditions, the capacity for rhizosphere acidification to increase Fe bioavailability may be insufficient to meet the elevated demand imposed by elevated [CO_2_] stimulated biomass accumulation. Additionally, Lindsay and Schwab (1982), suggest that redox potential (pe + pH) must stay below about 12 for enough Fe^2+^ to exist. Their results indicate the roots of Hawkeye (HA, PI 548577), a soybean variety, can lower the redox potential in surrounding soil by releasing electrons or other reducing agents into the root zone to increase soluble iron. Because the HA soybean line can acidify and chemically reduce the soil immediately around their roots (Lindsay and Schwab, 1982; Sain and Johnson, 1984), additional research could be performed to understand the soybean germplasm to better leverage the cation exchange. Incorporating these iron–efficient soybean lines as parents in breeding programs could offer a practical strategy to develop varieties with improved uptake potential and decreasing the need for external iron chelate applications.

The interaction between Fe and sulphur (S) metabolism is an additional consideration: Fe-S cluster assembly is a major sink for both elements in plant cells for (Balk and Pilon, 2011; Connorton *et al*., 2017). Fe is required for Fe–S clusters in photosynthetic and respiratory proteins, and S metabolism provides the sulphur needed for Fe–S cluster biosynthesis. Connorton *et al*. (2017) indicate that chloroplasts and mitochondria are major sites of Fe–S proteins and that cysteine provides sulphur for Fe–S cluster assembly. If elevated [CO_2_] changes S acquisition, N metabolism, or seed protein composition, Fe demand and Fe allocation may also shift. This does not mean Fe declines were caused by S limitation, but it provides a mechanistic basis for discussing why Fe may be more sensitive than nutrients whose uptake and allocation are less tightly linked to redox chemistry and cofactor assembly. Although S concentration declined under elevated [CO_2_], the broader nutrient-yield response suggests that soybean generally maintained macronutrient acquisition (**Fig. 5**; **Fig. 6**). However, because S metabolism is closely linked to protein synthesis and Fe–S cluster biosynthesis, changes in S availability or allocation could influence seed quality and Fe metabolism. Future work combining sulphate availability, root sulphur transporter, SULTR, expression, and Fe–S metabolic markers would help determine whether altered S acquisition contributes to Fe or protein declines under elevated [CO_2_].

### Hydraulic redistribution, nutrient stratification and nutrient recycling

A further mechanism that may modulate nutrient availability under elevated [CO_2_] is hydraulic redistribution – the passive movement of water from deep, moist soil layers into drier upper layers through root systems at night, driven by water potential gradients (Wan *et al*., 2000; Pang *et al*., 2013). This process can redistribute dissolved nutrients from deeper soil horizons to the upper root zone where active uptake predominantly occurs and may be altered under elevated [CO_2_] if root architecture or water use patterns change. HS, which showed greater stimulation of aboveground biomass under elevated [CO_2_] (**Fig. S2A & S2C**), may be particularly relevant in this context: greater carbon allocation to root systems with higher water use efficiency could modify the spatial distribution of nutrient depletion zones in the soil profile. If elevated [CO_2_] enhances hydraulic redistribution and increases nutrient mining from deeper layers, this could partly explain the observed compensatory uptake for many elements. Conversely, if greater belowground biomass under elevated [CO_2_] increases nutrient uptake from subsoil layers while surface soils become relatively depleted, this has implications for seasonal nutrient cycling and soil fertility management in soybean production systems.

The increase in whole-plant nutrient yield under elevated [CO_2_] observed in this study also has implications for nutrient cycling in cropping systems (**Fig. 6**). Nutrient yield in this context represents the total mass of mineral elements removed from the soil into plant tissue; a portion of this is exported from the field in harvested grain, while the remainder is returned to the soil in crop residues (leaves, stems, roots, and pod walls). The decomposition of this material by soil microorganisms constitutes the primary pathway by which nutrients fixed in plant tissues are remineralized and returned to the soil solution, where they become available for uptake by subsequent crops (McGrath *et al*., 2000; Sayer, 2006; Liu *et al*., 2022). Under elevated [CO_2_], increased aboveground biomass means that more nutrients are cycled through plant tissue each season, even if seed concentrations are reduced. The nutrients retained in residues that are not removed at harvest remain bioavailable for subsequent growing seasons. This is consistent with findings from forest ecosystems, where elevated [CO_2_] increased the total flux of N, P, K, Ca, Mg, and several micronutrients to the soil via litterfall, despite lower litter nutrient concentrations, because the stimulation of litter production exceeded the magnitude of concentration decline (Liu *et al*., 2007). However, litter produced under elevated [CO_2_] often has an elevated C:N ratio and higher lignin content, characteristics that reduce litter quality and can slow decomposition rates and delay nutrient release (Cotrufo *et al*., 1994; Norby *et al*., 2001). If decomposition rates of soybean residue are similarly reduced under elevated [CO_2_], the timing of nutrient release may be decoupled from crop demand in subsequent seasons, potentially reducing the agronomic value of residue-derived nutrients despite their greater total quantity. This trade-off between increased nutrient inputs via litter and potentially slower nutrient release rates under elevated [CO_2_], represents an important, and largely unresolved, question for the long-term management of soil fertility in soybean production systems under rising [CO_2_].

### Implications for human nutrition

Our finding that whole-plant and grain nutrient yield either increased or was maintained under elevated [CO_2_] for most elements measured here has direct bearing on the nutritional consequences of rising atmospheric CO_2_. Assessments of the health risks associated with elevated [CO₂] driven nutrient declines have largely focused on concentration data alone (Ainsworth *et al*., 2002; Loladze, 2014; Myers *et al*., 2014a) which implicitly assumes that yield gains are irrelevant to nutritional outcomes. Our data demonstrate that for most nutrients, including N, P, K, Ca, Mg and Zn, the yield stimulation under elevated [CO_2_] was sufficient to maintain or increase total nutrient delivery per unit land area, even in the face of concentration declines (**Fig. 6**). This does not resolve all questions about the net nutritional impact of elevated [CO_2_], downstream effects on food prices, dietary choices, and the broader nutritional context of affected populations remain important variables, but it substantially complicates the simple narrative that concentration declines necessarily translate to nutritional harm. The exception, again, is Fe: grain Fe yield declined under elevated [CO_2_], driven by both the strong concentration decline and the absence of a compensatory uptake response. Given the prevalence of Fe deficiency and soybean’s importance as a dietary Fe source in many regions, this finding underscores the need to better understand, and potentially engineer, the Fe acquisition pathway in soybean under future CO_2_ conditions.

## Conclusions

The decline in soybean mineral concentrations under elevated [CO_2_] reflects the combined influence of dilution, reduced transpiration-driven mass flow, and active root nutrient acquisition, with no single mechanism acting in isolation. Critically, active root uptake largely compensated for the negative effects of dilution and reduced mass flow for most nutrients, such that whole-plant and grain nutrient yield were maintained or increased under elevated [CO_2_] – a finding consistent with those of Digrado et al. (2024) and Kaur et al. (2025). These results challenge the assumption that mineral concentration declines under elevated [CO_2_] necessarily translate into reduced nutritional output from soybean. Iron was the sole exception, exhibiting consistent negative root uptake and the largest concentration decline of any element measured, likely reflecting pH-dependent constraints on Strategy I Fe acquisition that are not shared by other micronutrients. The two varieties differed substantially in yield response to elevated [CO_2_] but showed no difference in nutrient acquisition efficiency. These findings highlight the physiological plasticity of root nutrient acquisition as an underappreciated buffer against elevated [CO_2_] driven mineral dilution, while identifying Fe as a nutritionally significant vulnerability that warrants targeted attention in breeding and soil management strategies under future atmospheric CO_2_ conditions.

## Acknowledgements

We thank Jesse McGrath, Christopher M. Montes, and Eric Vargas for setting up the CO_2_ fumigation system and ensuring the experiment ran smoothly; Anthony Digrado and C.M.M for planting in all growing seasons; and Isabelle Gawedski, Nicole Miedzinski, Lilah Ranz, Ximin Piao, and Eneji D. Sani for helping to install sapflow sensors and collect field data. Special thanks to Elizabeth Ainsworth, Carl Bernacci, Emily Heaton, Christopher M. Montes, and Scott Oswald for helpful discussions.

We acknowledge the USDA ARS and the University of Illinois at Urbana-Champaign for funding SoyFACE. Any opinions, findings, and conclusions or recommendations expressed in this publication are those of the authors and do not necessarily reflect the views of the US Department of Agriculture. Mention of trade names or commercial products in this publication is solely for the purpose of providing specific information and does not imply recommendation or endorsement by the US Department of Agriculture. USDA is an equal opportunity provider and employer.

## Author contributions

Terence Seldon Kwafo: conceptualization, modelling, data curation; formal analysis; methodology; visualization; writing – original draft; writing – review and editing

Kylie Yerkes: data curation, data organisation, and data analysis

Justin M. McGrath: conceptualization; modelling; funding acquisition; investigation; project administration; resources; supervision; review and editing

## Data availability

Data for reproducing all analyses contained in this work are available upon request.

