## Supplemental data for "Nutrient Concentration Declines but Nutrient Yield Increases Under Elevated CO_2_ Concentration in Soybean: Disentangling the relative effects of Dilution, Transpiration, and Uptake Activity"

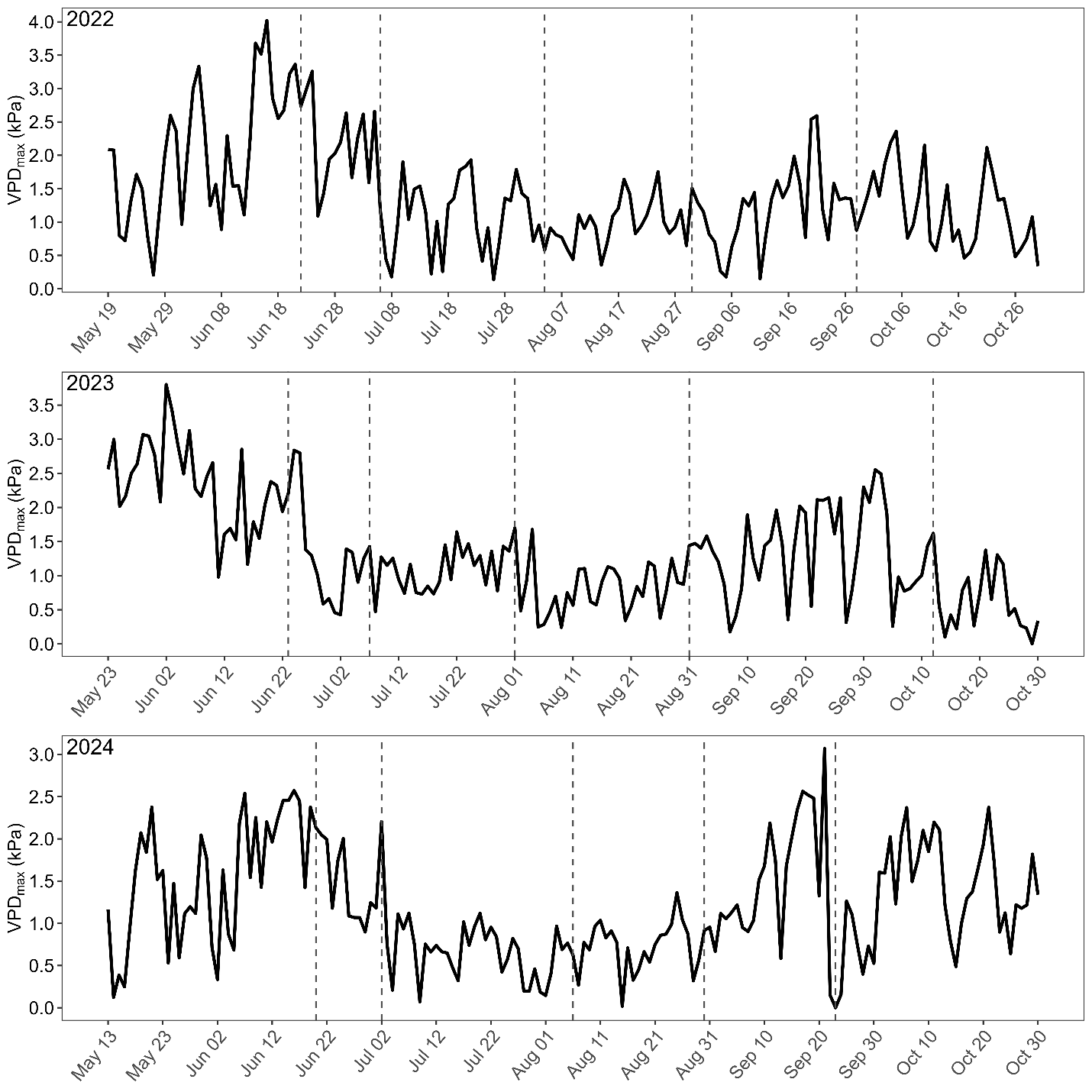

**Fig. S1.** Maximum vapour pressure deficit (VPD_max_). Data were obtained from SoyFACE weather station. Dash vertical lines denote sampling dates.

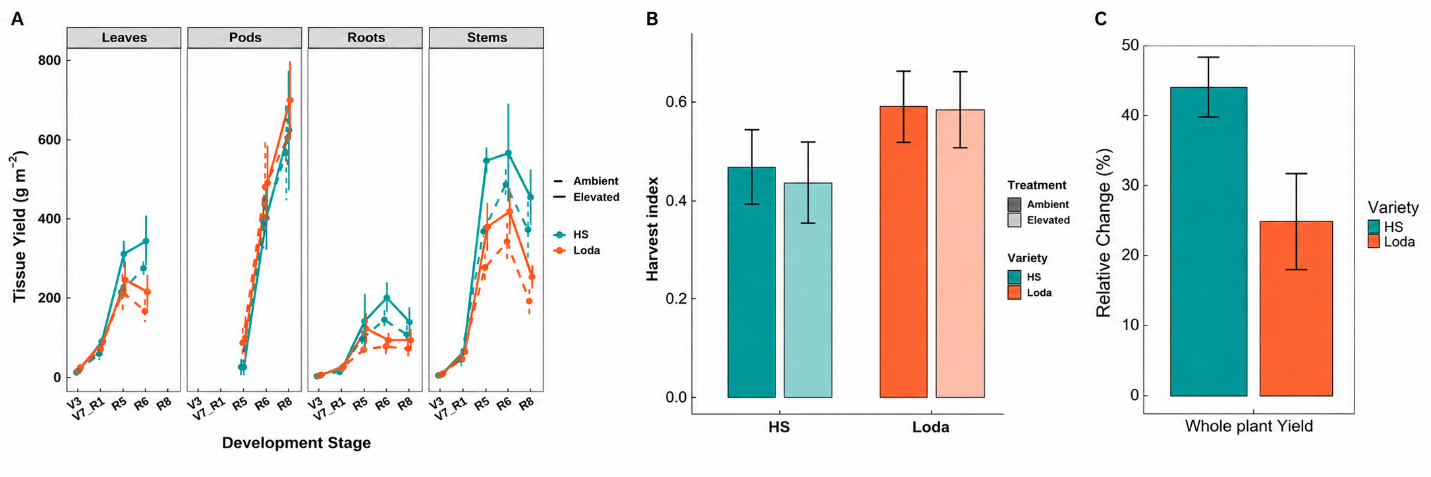

**Fig. S2. (A)** Partitioned tissue yield. **(B)** Harvest Index. (**C**) Change (%) in whole plant biomass yield response to elevated CO_2_ relative to ambient at seed fill (R5). HS93-4118 (HS) and Loda are represented in blue and orange, respectively. Error bars represent 95% CI.

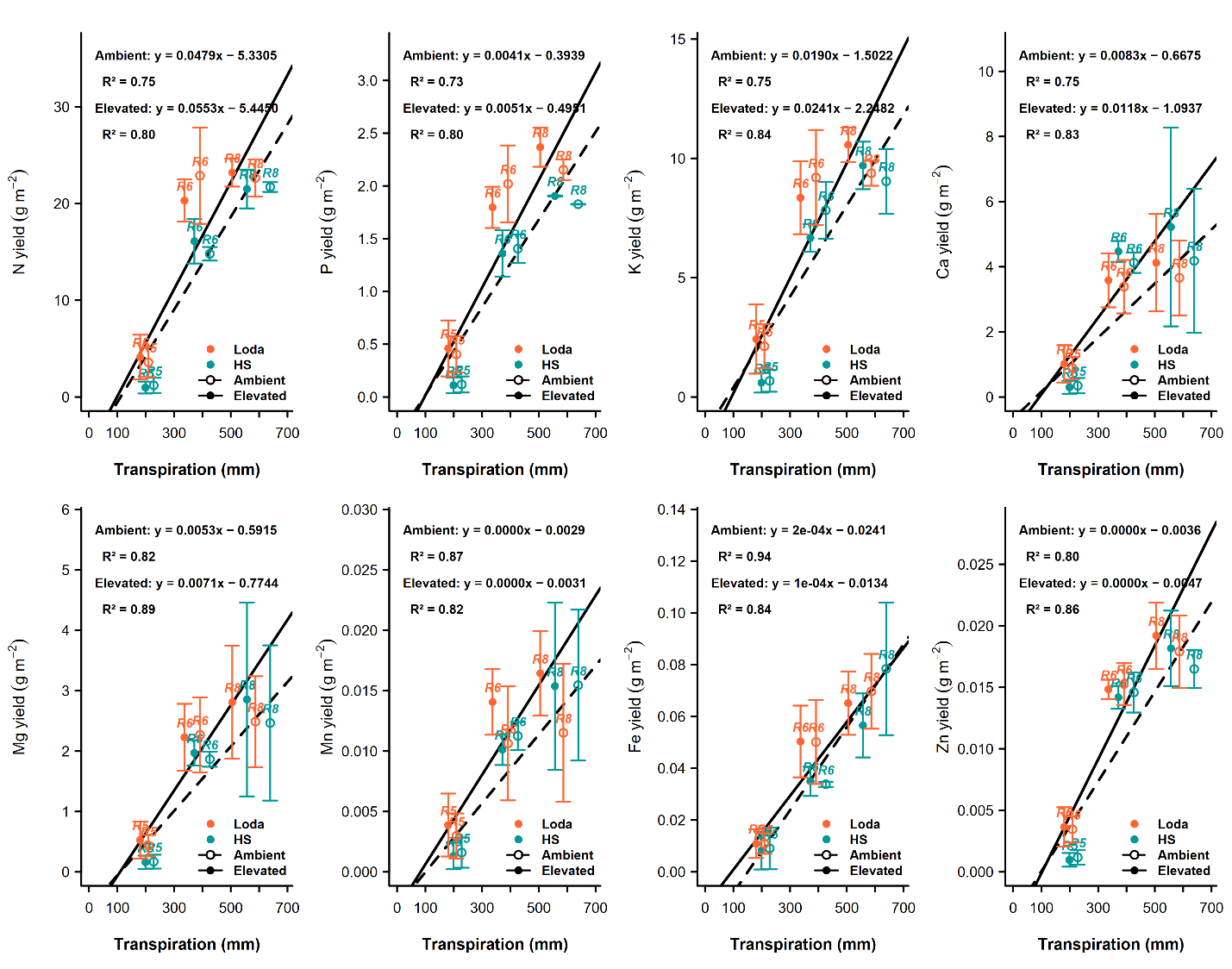

**Fig. S3.** **Effects of transpiration on nutrient uptake in response to elevated CO_2_.** Grain nutrient response at final harvest (R8)**.** HS93-4118 (HS) and Loda are represented in blue and orange, respectively. Error bars represent 95% CI.

**Table S1. Dates of planting, harvest, and sampling.**

| **Year** | **Planting date** | **V3** | **V7/R1** | **R5** | **R6** | **R8 (Final harvest)** |
| --- | --- | --- | --- | --- | --- | --- |
| 2022 | May-19 | Jun-22 | Jul-06 | Aug-04 | Aug-30 | Sep-28 |
| 2023 | May-23 | Jun-23 | Jul-07 | Aug-01 | Aug-31 | Oct-12 |
| 2024 | May-13 | Jun-20 | Jul-02 | Aug-06 | Aug-30 | Oct-12 |

**Table S2. Total precipitation, average maximum air temperature, and average maximum VPD from 2022 to 2024.**

| **Year** | **Total precipitation (mm)** | **Average maximum air temperature (℃)** | **Average maximum VPD (kPa)** |
| --- | --- | --- | --- |
| 2022 | 235 | 27.9 | 1.46 |
| 2023 | 229 | 27.6 | 1.42 |
| 2024 | 475 | 27.3 | 1.17 |

**Table S3. Soil and soil water nutrient composition. All soil nutrients are in ppm (mg/kg).**

| **(A) Soil** | | | | | | | | | | | | | |
| --- | --- | --- | --- | --- | --- | --- | --- | --- | --- | --- | --- | --- | --- |
| Year | pH | OM (%) | CE | Nitrate | P | K | Ca | Mg | S | Zn | Mn | Fe | Cu |
|  |  |  | (meq/100g) |  |  |  |  |  |  |  |  |  |  |
| 2022 | 6.1 | 3.9 | 19.9 |  | 11.0 | 112.0 | 2470.0 | 514.0 | 8.0 | 1.1 | 28.0 | 108.0 | 1.5 |
| 2023 | 6.4 | 4.8 | 17.6 | 8.6 | 20.0 | 116.0 | 2402.0 | 455.0 | 2.8 | 2.7 | 19.0 | 113.0 | 1.8 |
| 2024 | 6.8 | 1.9 | 9.5 | 3.1 | 9.0 | 72.5 | 1372.0 | 253.0 | 1.0 | 0.6 | 15.0 | 81.0 | 0.9 |

| **(B) Soil water** | | | | | | | | | | | | |
| --- | --- | --- | --- | --- | --- | --- | --- | --- | --- | --- | --- | --- |
| Year | pH | Ecw | Nitrate | P | K | Ca | Mg | S | Zn | Mn | Fe | Cu |
| 2022 | 7.8 | 0.44 | 27.8 | 0.021 | 2.42 | 51.4 | 20.4 | 11.7 | 0.039 | 0.0083 | 0.079 | 0.0044 |
| 2023 | 7.3 | 0.38 | 51.3 | 0.16 | 1.9 | 46.3 | 14.2 | 11.4 | 0.062 | 0.0115 | 0.1 | 0.005 |
| 2024 | 7.6 | 0.36 | 20.45 | 0.16 | 1.6 | 42.1 | 16.1 | 10.7 | 0.05 | 0.0111 | 0.1 | 0.005 |
| **Mean** | **7.6** | **0.39** | **33.2** | **0.11** | **1.97** | **46.6** | **16.9** | **11.3** | **0.050** | **0.0103** | **0.093** | **0.0048** |

Equations:

$$\begin{aligned} \text{r=}\frac{\text{(}\text{B}_{\text{elev}}\text{-}\text{B}_{\text{amb}}\text{)}}{\text{B}_{\text{amb}}}\text{=}\frac{\text{B}_{\text{elev}}}{\text{B}_{\text{amb}}}-1 \#\left( 1 \right) \end{aligned}$$

Dilution only effect, Concentration = (ambient yield of nutrient X) / (elevated biomass)

$$\text{[X]}_{\text{elev}}\text{=}\frac{\text{Y}_{\text{amb}}}{\text{B}_{\text{elev}}}\text{=}\frac{\text{Y}_{\text{amb}}}{\text{B}_{\text{amb}}\text{⋅}\text{(1+r)}}$$

The percent change in concentration relative to ambient would then be:

$$\begin{aligned} \frac{\left[ \text{X}\text{]}_{\text{elev}}\text{ - [X} \right]_{\text{amb}}}{\left[ \text{X} \right]_{\text{amb}}}\text{×100=}\frac{\frac{\text{Y}_{\text{amb}}}{\text{B}_{\text{amb}}\text{(1+r)}}\text{-}\frac{\text{Y}_{\text{amb}}}{\text{B}_{\text{amb}}}}{\frac{\text{Y}_{\text{amb}}}{\text{B}_{\text{amb}}}}\text{×100=}\frac{\text{-r}}{\text{1+r}}\text{×100} \#\left( 2 \right) \end{aligned}$$

The observed change in concentration integrates all of the above mechanisms and was calculated directly from measured tissue concentrations at ambient (s₀) and elevated (s₁) CO_2_:

$$\begin{aligned} \text{Observed }\Delta\text{conc}=\frac{s_{1}-s_{0}}{s_{0}}\times100 \#\left( 3 \right) \end{aligned}$$
